# Monocyte Infection Dynamics Shape HIV-1 Phyloanatomy in the Peripheral Blood

**DOI:** 10.1101/494856

**Authors:** Brittany Rife Magalis, Samantha L. Strickland, Stephen D. Shank, Patrick Autissier, Alexandra Schuetz, Pasiri Sithinamsuwan, Sukalaya Lerdlum, James L. K. Fletcher, Mark de Souza, Jintanat Ananworanich, Victor Valcour, with the SEARCH007 Study Group, Nicholas Hatchings, Yuwadee Phuang-Ngern, Weerawan Chuenarom, Somporn Tipsuk, Mantana Pothisri, Tanate Jadwattanakul, Supun-nee Jirajariyavej, Chayada Sajjaweerawan, Boot Kaewboon, Siriwat Akapirat, Thep Chalermchai, Duanghathai Suttichom, Peeriya Prueksakaew, Putthachard Karnsomlap, David Clifford, Robert H. Paul, Jerome H. Kim, Kenneth C. Williams, Sergei L Kosakovsky Pond, Silvia Ratto Kim, Marco Salemi

## Abstract

Human immunodeficiency virus (HIV) RNA and DNA have been isolated from patient monocytes, an immune cell population that is quite different in several aspects from the canonical T-cell viral target. Because monocytes are migratory and resilient to both natural and synthetic antiviral defenses, knowledge of the contribution of monocyte infection to ongoing viral evolution and spread *in vivo* is of significant interest for drug development and treatment strategies. Using single viral genome sequencing from different peripheral blood compartments and phyloanatomic statistical inference, we demonstrate that productively infected monocytes follow an evolutionary trajectory that is distinct from peripheral T cells during multiple stages of disease progression. Gene flow and selection analysis reveal plasticity in the source of monocyte infection and in the region of the HIV envelope glycoprotein that experiences selection pressure across individuals. The findings, thus, point to a potential reservoir showing a range of infection and transmission dynamics, for which the current universal, T cell-targeted treatment strategies would be inadequate.

**Author summary:** Human immunodeficiency virus is a rapidly evolving virus, allowing its genetic material to act as a fingerprint for epidemiological processes among, as well as within, individual infected hosts. In this study, sampling of viral RNA from plasma and differing infected immune cell populations from the peripheral blood was undertaken for three separate individuals in order to infer such processes. The results revealed a productively infected monocyte cell population, for which distinct selection pressures were linked to differing spatiotemporal origins of infection. In light of previous evidence of the migratory nature of these cells and resilience to both natural and synthetic antiviral defenses, this study underscores the importance of further investigation into the role of monocytes, and monocyte-rich tissues, as viral reservoirs during treatment of HIV with antiretroviral therapy.

## Introduction

The primary target for human immunodeficiency virus (HIV) infection is CD4+ T cells, though the virus has been detected in virtually every tissue and organ system within the human host [1]. Macrophages found within HIV-infected tissues are an important additional cellular target for the virus, with relevance to transmission, persistence in the presence of antiretroviral therapy (ART), and development of AIDS-related comorbidities [2], particularly neurocognitive impairment. [3, 4] Various types of tissue-resident macrophages originate from circulating blood monocytes, which are recruited at a high rate to sites of inflammation during HIV infection [5]. In addition to acting as macrophage precursors, recent studies report that monocytes function to constitutively traffic captured antigen between tissues and corresponding draining lymph nodes. [6, 7]. Unlike T cells, however, monocytes (and macrophages) are relatively resistant to the cytopathic effects of the virus. Therefore, in the face of ART, monocytes represent a potentially significant reservoir, capable of disseminating virus across a vast anatomical space for longer periods of time and to anatomical viral sanctuaries wherein ART penetration is poor. [8, 9]

The low frequency (<1%) of HIV-infected monocytes in the peripheral blood and the difficulty of obtaining tissue samples present major barriers to elucidating the dynamics of HIV infection in human monocytes. [10] Earlier studies focusing on circulating monocytes revealed genetic patterns distinct from that of T cells, [11–13] consistent with the hypothesis of differing viral population dynamics. However, investigation of the role(s) of monocytes in the evolution and spread of HIV-1 has been largely limited to the differentiated macrophage phenotype. [14] Difficulties in isolating viral genomic RNA, indicative of replicating virus, from patient-derived monocytes [15] have also impeded the inclusion of this cell population in models of evolutionary trajectories among differing anatomical compartments. [16–19] In this study we demonstrate the successful isolation of monocyte viral RNA for use in a phyloanatomic study [16] of the evolutionary dynamics and genetic features of replicating HIV in the peripheral blood of three subtype CRF01_AE-infected individuals. The findings of this study reveal a variable role for circulating monocytes in HIV gene flow within the blood, with putative links to region-specific selection pressures within the envelope glycoprotein gene.

## Materials and methods

### Ethics Statement

Fifty HIV-1-infected, cART-naive volunteers were enrolled into the SEARCH 007 study (registration number NCT00777426) at the Thai Red Cross AIDS Research Center in Bangkok, Thailand [20]. All volunteers met the Thai Ministry of Health guidelines to initiate therapy based on having symptomatic HIV infection or a CD4+ T-cell count below 350 cells/mL [21]. Signed informed consent was obtained for all participants, consisting of adults at least 18 years of age. The Chulalongkorn University Institutional Review Board in Bangkok, Thailand, and the University of Hawaii approved the study. Details of the clinical trial protocol can be found in Supplementary Information.

### Enrollment of study population

Cognitively normal (NL) subjects were matched to NCI individuals by age (within a decade), education (less than a high school degree, high school degree, some college, college degree), gender, and CD4+ T-cell count. In addition, 10 HIV-uninfected controls (CL) were enrolled. HIV-infected volunteers were categorized as NL or having NCI, with nine volunteers meeting criteria for Asymptomatic Neurocognitive Impairment (ANI), nine – for Mild Neurocognitive Disorder (MND), and nine having HIV-Associated Dementia (HAD) based on a clinical assessment/ neuropsychological testing. Clinical examination included a neuropsychological testing battery [22], the International HIV Dementia Scale [23], a macroneurological examination, and brain magnetic resonance imaging/spectroscopy (MRI/MRS). Final diagnoses were assigned by consensus conference that included an HIV neurologist and neuropsychologist using established criteria [24]. HIV-uninfected participants were selected using the same inclusion and exclusion criteria, with the exception of CD4+ T-cell count. These exclusion criteria encompassed previous exposure to antiretroviral therapy, positive Hepatitis C serology, and presence of factors that could cause cognitive abnormalities (e.g., past head injury, learning disabilities, major depression, illicit drug use, active opportunistic infection, past or current CNS infection). Individuals were required to have a negative urine drug test prior to enrollment. Lumbar puncture and brain MRI were performed, if indicated, to exclude central nervous system (CNS) opportunistic infection. HIV+ volunteers initiated therapy (zidovudine or stavudine, lamivudine, and nevirapine) at time of enrollment and returned to the clinic at three-month intervals for the duration of one year. Peripheral mononuclear cells (PBMCs) were collected and neuropsychological testing performed at baseline and twelve months following enrollment/cART initiation. Data were generated from baseline and at twelve months post-cART initiation. Groups were analyzed based on NCI diagnosis at the time of baseline assessment [25].

### Cell sorting

Cryopreserved PBMCs were quickly thawed in a 37°C water bath before being transferred to a 50ml conical tube containing 40ml RPMI with 20% FBS pre-warmed at 37°C. Cells were washed twice and transferred to a FACS tube and stained for 15 minutes at room temperature with an antibody cocktail consisting of anti-CD14-Pacific Blue (clone M5E2), anti-CD3-Alexa Fluor 700 (clone SP34-2), anti-CD20-Cy7-APC (clone B27) and anti-CD16-Cy7-PE (clone 3G8) (all from BD Pharmingen, San Jose, CA), anti-HLA-DR-ECD (clone L243, Beckman Coulter, Miami, FL), and Live/Dead Aqua (Invitrogen, Eugene, OR). All antibodies were titrated to determine optimal concentrations. Antibody-capture beads (CompBeads, BD Biosciences) were used for single-color compensation controls for each reagent used in the study, with the exception of cells being used for anti-CD3 and Live/Dead Aqua. After staining, cells were washed once, filtered and resuspended in 1ml PBS. The BD FACSAria cytometer (BD Biosciences, San Jose, CA) was set up with a pressure of 20 psi and a 100-um nozzle was used. Instrument calibration was checked daily by use of rainbow fluorescent particles (BD Biosciences). After acquiring unstained and single-color control samples to calculate the compensation matrix, we acquired 1 x 106 events in order to set up the sorting gating strategy. CD14+ monocyte population were gated first based on FSC and SSC parameters, after which we excluded 1) dead cells by gating out Aqua+ cells and 2) unwanted cells by gating out CD3+ and CD20+ cells and then gated on HLA-DR+ cells. From the HLA-DR+ population, a dot plot of CD14 vs. CD16 was used to make a sorting gate, which included all monocytes except the CD14-CD16-subset. For CD3+ T-lymphocyte sorting, FSC and SSC parameters were used to gate lymphocytes, dead cells were excluded by using Aqua staining, and CD14+ cells were also excluded. Following this procedure, the CD3+ T-lymphocytes were gated based on CD3 expression and negativity for CD16. Post-sort purity were checked for each sample, and both CD14+ and CD3+ sorted subpopulations were >98% pure. After cell sorting, the highly pure cell populations were washed with PBS twice and all liquid was aspirated. Cells were then stored as dry pellets at −80°C.

### RNA extraction and cDNA synthesis

Cell-free viral RNA was extracted from participant plasma using the Qiagen QIAamp Viral RNA Mini Kit, whereas sorted PBMC-associated RNA and DNA were processed using the Qiagen Allprep DNA/RNA Mini kit according to the manufacturer’s protocols. Viral RNA was then reverse transcribed into cDNA according to the manufacturer’s protocol using the SuperScript®III First-Strand Synthesis System kit (Invitrogen). The following primer was used for reverse transcription: ’K-env-R1’ 5’-CCAATCAGGGAAGAAGCCTTG-3’ (HXB2 coordinates 8736-8716) [26].

### Single genome amplification and sequencing

HIV-1 env *gp120* sequences were amplified from viral cDNA and genomic DNA (gDNA) using a modified limiting-dilution two-round PCR approach (‘single genome sequencing’) based on previously published methods [27] in order to prevent PCR-mediated resampling and recombination. The following primers were used for both rounds of PCR: ’polenv_AE’ 5’-GAGCAGAAGACAGTGGAAATGA-3’ (HXB2 coordinates 6207-6228; modified from Tuttle et al., 2002 [28] for subtype AE) and ’192H’ 5’-CCATAGTGCTTCCTGCTGCT-3’ (HXB2 coordinates 7815-7796; modified from Maureen Goodenow for subtype AE). PCR reactions consisted of 2 minutes at 94°C for 1 cycle, 30 seconds at 94° C, 30 seconds at 58°C, and 3 minutes at 72°C for 40 cycles, then 10 minutes at 72°C using the Platinum®Blue PCR SuperMix (Invitrogen). Amplicons were then visualized using 1% agarose gel electrophoresis with an Amplisize^TM^ Molecular Ruler 50-2,000 base pair (bp) ladder (Bio-Rad). Sequencing was performed using an Applied Biosystems 3730xl DNA Analyzer (Life Technologies) at the University of Florida Interdisciplinary Center for Biotechnology Research genomics core facility.

RNA and DNA extractions, cDNA synthesis and first round PCR set-up were conducted in a restricted-access amplicon-free room with separate air-handling, with laboratory equipment where no amplified PCR products or recombinant cloned plasmids were allowed and where work surfaces and equipment were thoroughly cleaned before and after use with Eliminase®(Decon Labs, Inc.). PCR loading was performed so as to minimize contamination across plasma and cell-specific samples for individual participants. For example, PCR amplification plate #162 contained diluted RNA from P01V1 and P02V1, but only for monocytes.

### Sequence alignment and analysis

Individual nucleotide sequence chromatograms were visualized using Geneious vR6 [29] for the investigation of sites assigned multiple nucleotide identities and for removal of potential PCR errors identified as singleton insertions or deletions. Nucleotide changes present in ≤ 1% per site were reverted to the nucleotide with the highest frequency [30]. Sequences are available in GenBank (XXXX-XXXX). Patient-specific sequences were translated and aligned using the Clustal algorithm [31] implemented in BioEdit v7.1.11 [32] (available from http://www.mbio.ncsu.edu/bioedit/bioedit.html) followed by manual optimization of positional homology [33] and removal of gap-filled regions within the hypervariable V1V2 domains. The final alignment included 1,068 nucleotides spanning position 6381-7580 of the HXB2 reference strain. Putative intra-host recombinants were identified using SplitsTree4 software [34] and removed prior to phylogenetic analysis. Alignments are available from GitHub (see Data Availability). Neighbor-joining (NJ) tree reconstruction was then performed using MEGA v5.2.2 [35] with the HKY model of nucleotide substitution [36] and gamma-distributed rate variation across sites. Pairwise deletion was used for treatment of gaps within the alignment. Branch support was assessed by bootstrapping (1,000 replicates). Sequences from all participants were included in the NJ tree in order to infer participant viral subtype and the extent of sequencing cross-contamination based on participant-specific clustering patterns (S10 Fig).

Evolutionary analysis was performed for participants from whom a sufficient number of monocyte-derived sequences were available to produce adequate phylogenetic signal for the monocyte compartment (P01, P02, and P13). Sequences from two separate time points (0 and 12 months) were analyzed for P01, who maintained MND diagnosis throughout the study and did not suppress viral load, despite the initiation of cART upon enrollment. Viral genetic diversity was quantified by pairwise genetic distances estimated for sequences derived from cell-free virus in the plasma and from sorted peripheral CD3+ T-lymphocytes and CD14+ monocytes within the three previously described individuals (P01, P02, and 013). This estimation was performed in R (APE package) [37] using the TN93 nucleotide substitution model [38] with gamma distributed rate variation across sites (α=0.1). A viral epidemiology signature pattern analysis (VESPA) was used to detect distinct frequency variation in particular amino acids between plasma, CD3+ T-lymphocyte, and CD14+ monocyte viral sequences for P01, P02, and P13.

### Maximum likelihood tree reconstruction and compartmentalization analysis

Because viral population structure, such as that dependent on anatomical location or cell type, can affect patterns of polymorphism that contribute to significant genetic variation or that mimic selection [39], the extent of this structure was assessed both qualitatively and quantitatively within each participant. Maximum likelihood (ML) tree reconstruction was performed for each of the three individuals using all available *gp120* sequences in order to assess clustering patterns according to anatomical location and time of sampling and was performed in IQTree [40] using the best-fit evolutionary model according to the Bayesian Information Criterion. Tree correlation coefficients (TCC) were estimated to provide a quantitative assessment of compartmentalization, representing the relationship between population isolation and the distance within the tree, with population subdivision defined in this study in terms of either space or time and tree distance measured according to the number of branches (*r*_*b*_) or patristic distances (*r*) separating two sequences [41, 42]. Anatomical compartmentalized structure was also analyzed in order to determine if within-host epidemiological linkages between peripheral cell populations and plasma could be resolved reliably. Clustering patterns within the ML and Bayesian (see below) phylogenies based on anatomical sampling origin, particularly a mixture of paraphyletic and polyphyletic clades, were used to exclude the possibility of significant intermediary viral subpopulations or common sources of virus when interpreting the results of the Bayesian phyloanatomy analysis in BEAST [43].

### Selection Analysis

Selection analyses were performed using a modification of the Fixed Effects Likelihood (FEL) for estimating site-specific selective pressures [44]. The test, referred to as contrast-FEL (or cFEL), is available in HYPHY [45], v2.3.14 or later. Briefly, each intra-host ML tree containing all available sequences for each individual was partitioned into groups of branches according to their compartment membership (internal branches were labeled with a compartment if and only if all of their descendants belonged to the same compartment), generating 4 sets of branches: Plasma, CD3, CD14, and background (i.e. not labeled). We next fitted the MG94xREV model to the entire alignment by maximum likelihood to estimate nucleotide substitution biases and relative branch lengths. Finally, for every site, we fitted a model with 5 parameters: *α* (site-wide synonymous rate, relative to the gene average), and four compartment-specific non-synonymous substitution rates *β*_*K*_, where *K ∈* Plasma, CD3, CD14, background. Annotated trees and alignments for patients P01, P02, and P13, along with cFEL results and mapped cFEL-identified sites on the 3D Env structure (3JWO) [46]), are available at http://thai.hyphy.org/ and exemplified in Fig 1.

**Figure 1.**
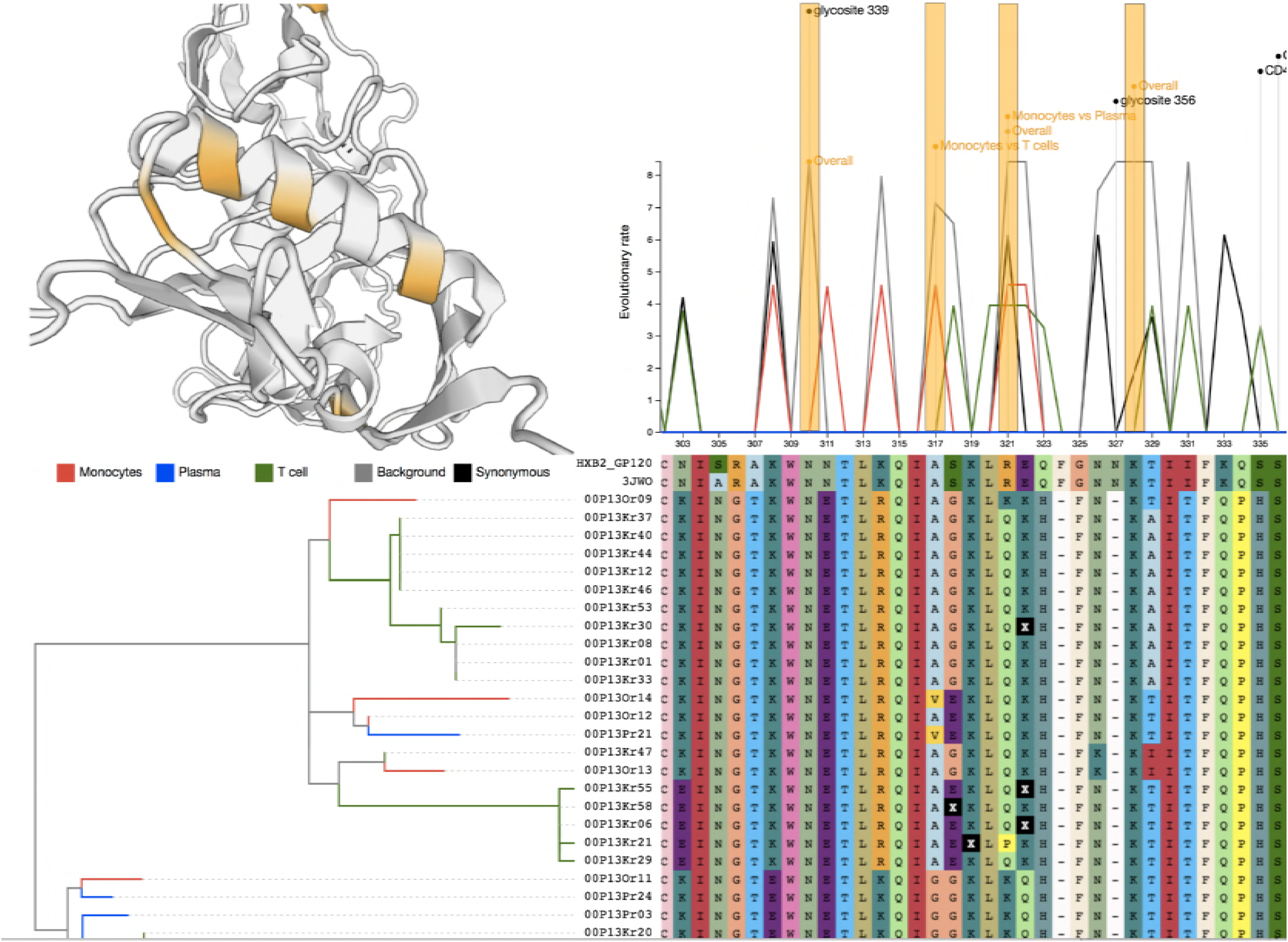
Visualization tool for the results of contrast-fixed effects likelihood (cFEL) estimation of site-specific selection pressure in the context of the alignment, tree, and protein 3D structure. Results are pictured for P13 of this study. The maximum likelihood tree (bottom left) is annotated according to anatomical grouping. The alignment (bottom right) is annotated (top right) according to Los Alamos National Laboratory HIV Database HXB2 sequence annotation https://www.hiv.lanl.gov/content/sequence/HIV/MAP/annotation.html. The 3D structure for Env (3JWO, top left) was determined by Pancera et al. [46]). The cFEL results panel (also top right) indicates normalized (by branch length) rates of synonymous (black) and non-synonymous (color according to group) substitutions within the tree. Sites experiencing significant differences in selection among groups have been highlighted in orange and mapped to the 3D structure. Interactive visualization of the results for patients P01, P02, and P13 can be found at http://thai.hyphy.org/.

### Bayesian phyloanatomic reconstruction

Because sampling was not uniform across sampling locations for these three participants, three replicates of random sampling (without replacement) according to the minimal number of available sequences in one of the three locations (P001=15, P002=20, P013=10) was performed for each of the three participants (with equal representation of months 0 and 12 for P01) in order to reduce the impact of spatial sampling bias while still incorporating the information from all sequences in the Bayesian phylogenetic analysis. Phylogenetic signal was determined prior to Bayesian analysis for each of the participant-specific sample replicates using likelihood mapping [47] implemented in TreePuzzle v5.2 [48] (available from http://www.tree-puzzle.de/), the results of which indicated sufficient signal for phylogenetic analysis (S11 Fig)). Temporal signal was also assessed for individual sample replicates of P01, consisting of sequences sampled at multiple time points, by determining the significance of the relationship of sequence sampling time to genetic divergence from the most recent common ancestor of all sequences within the transmission cluster. We used a clustered permutation approach in BEAST [49, 50] (available from http://beast.bio.ed.ac.uk/), asking whether the correlation was stronger than expected if sampling dates were randomly assigned among clusters of sequences sampled on the same date [51]. Clustered permutation tip date randomization [51, 52] was performed in R using the TipDatingBeast package for 5 replicates [53] and used for Bayesian evolutionary reconstruction in BEAST assuming the uncorrelated, relaxed molecular clock [54] and Bayesian skyline demographic models [50]. Markov chain Monte Carlo sampling of parameters and tree topologies was performed for 500 million generations or until effective sample sizes (ESS) reached values greater than 200 (after burn-in of 10%). ESS were calculated in Tracer (available from http://beast.bio.ed.ac.uk/Tracer). The Bayesian stochastic search variable selection model (BSSVS) [17] of asymmetric transition rates among discrete anatomical locations was incorporated into the non-randomized tip date BEAST analysis, as migration rates are assumed to be independent of evolutionary reconstruction (i.e., no impact on dating). Using an asymmetric transition rate matrix within the BSSVS model allowed for inferred directionality of significantly non-zero rates of viral dispersion between sampled anatomical compartments (plasma, CD3+ T-lymphocytes, and CD14+ monocytes). The hierarchical phylogenetic model was used to summarize trends across the three sample replicates for each participant [55]. As mentioned previously, because monocyte sequences were unobtainable at later sampling time points for P001 and P002, only sequences from the first visit (V1) for these two participants were used for phyloanatomic analysis. Given the robustness of the molecular clock for contemporaneously sampled HIV sequences [56], relaxed clock calibration was enforced with a mean evolutionary rate of 6.82E-04 substitutions/site/month, based on previous estimates [57]. Detailed information regarding additional evolutionary parameters and associated priors used in BEAST analysis is available upon request. Trees sampled (1, 000) from the posterior distribution (after burn-in of 10%) were visualized simultaneously and branch density assessed using DensiTree (available from https://www.cs.auckland.ac.nz/∼remco/DensiTree/), with high-density areas indicative of increased certainty of clustering patterns.

### Statistical Analysis

Significant differences for compartment-specific viral diversity (pairwise genetic distance) between participants were determined using a non-parametric pairwise multiple comparisons analysis based on rank sums (Dunn test package in R) with Bonferroni p-value correction following rejection of the D’Agostino Pearson test of normality. A p-value of ≤0.5 was considered significant. Statistical significance for Critchlow’s correlation coefficient [58] was determined using a null distribution of permutated sequences (1,000 permutations). A p-value of ≥0.5 was considered significant. For the tip date randomization test, absence of overlap between 95% high posterior density intervals of the mean evolutionary rate of the randomized and the correctly dated run indicated significant temporal signal (S8 Fig). With regard to the BSSVS analysis, Bayes Factors (BF) were calculated according to Lemey et al. (2009) [17] for participant-specific transition rates between each compartment using the MCMC rate posterior odds output from BEAST; BSSVS transition rates with BF>3 were considered to be well supported [17]. To test for differences in selection among compartments, we created null models by either constraining all non-synonymous rates to be equal, or by constraining pairs of non-background rates to be equal. p-values were derived using the likelihood ratio test assumed to follow the asymptotic 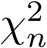 statistic (*n* = 3 for the “all” test, or *n* = 1 for pairwise tests). The Holm-Bonferroni procedure was employed for multiple test correction at each site (4 tests). Corrected p-values of 0.05 or less were considered significant.

### Data Availability

The authors declare that all data supporting the findings of this study are available within the paper (and its Supplementary Information files), but original data that supports the findings are available from GenBank (XXXX-XXXX) and http://http://thai.hyphy.org/, or the corresponding authors upon reasonable request.

## Results

Peripheral blood samples were obtained from individuals (CD4 count <350 cells/mm^3^) enrolled in a volunteer longitudinal HIV study in Thailand. [20] Using sorting techniques relying on immune-specific cluster of differentiation (CD) cell surface markers, we isolated, with ≥98% purity, monocyte (CD14+) and T cell (CD3+) populations (S1 Fig) in thirteen of twenty-two individuals (Table 1). We generated full-length envelope glycoprotein *gp120* monocyte-derived sequences in ten of the participants using single genome sequencing (SGS [27], S2 Fig). Though evidence of productive infection was only present in a subset of individuals, infection in the remaining individuals at levels below the assay limit of detection cannot be ruled out. Higher viral loads (VL) were positively correlated with the normalized number of RNA genome copies (*R*^2^ = 0.4, S3 Fig) and were marginally predictive of the ability to successfully sequence viral RNA (*p* = 0.06, logistic regression, S4 Fig). In three of the participants (P01, P02, and P13) with VL> 10^5^ copies/mL (Table 1 and S4 Fig), we successfully generated 10-48 *gp120* sequences from each of the peripheral monocyte, T-cell, and plasma compartments, permitting meaningful phylogenetic inference. Patient P01 was not successfully virally suppressed 12 months following initiation of combination antiretroviral therapy (cART) and sampling, suggesting non-adherence to therapy, though time of interruption is unknown.

**Table 1.** Viral burden in plasma and sorted PBMCs among Thai cohort.

| Participant | Diagnosis | Time since 1st visit (cART initiation) |  |  |  |  |  |
| --- | --- | --- | --- | --- | --- | --- | --- |
|  |  | 0 months (V1) |  |  | 12 months (V5) |  |  |
|  |  | Viral Load<br>(copies/mL) | CD14+<br>HIV RNA<br>Genomes# | SE | Viral Load<br>(copies/mL) | CD14+<br>HIV RNA<br>Genomes# | SE |
| **P002\*** | NL | 397150 | 156548 | 30127 | 263 | ND | - |
| P024 | NL | 232930 | 5314 | 1811 | <50 | ND | - |
| P026 | NL | 99649 | ND | - | <50 | ND | - |
| **P019** | NL | 98808 | 4037 | 2663 | <50 | ND | - |
| **P003** | ANI | 96396 | 6773 | 2709 | <50 | ND | - |
| **P007** | ANI | 46885 | 3434 | 1919 | <50 | ND | - |
| P008 | ANI | 16616 | 1677 | 1572 | <50 | ND | - |
| **P001\*** | MND | 750000 | 269875 | 27639 | 100000 | 129484 | 36435 |
| **P013\*** | HAD | 540909 | 12038 | 3398 | <50 | ND | - |
| **P015** | HAD | 350439 | 5422 | 3030 | <50 | ND | - |
| **P021** | HAD | 385478 | 4698 | 2181 | <50 | ND | - |
| **P004** | HAD | 183154 | 24680 | 10212 | <50 | ND | - |
| **P029** | HAD | 17277 | 6793 | 2445 | <50 | ND | - |

\*Copies per million sorted CD3+ (T cell) or CD14+ (monocyte) PBMCs and standard error (SE) as estimated in QUALITY [59] using limiting dilution PCR results, unless not detectable (ND).
\*Participants from which HIV genomic RNA was PCR-amplified (bold) and 10 or more sequences per cell type were successfully obtained.

We observed significant differences in pairwise sequence diversity (%) between monocyte-derived *gp120* sequences and those of plasma and T cells (S6 Fig), in agreement with earlier studies that used viral DNA [12, 60]. Patterns of diversity differed among patients and sampling times. In P02, HIV isolates sampled from monocytes (2.1%) were significantly less diverse (*p* < 0.001) than those from plasma (3.2%) and T cells (5.0%). In P13, monocyte HIV diversity (2.4%) was significantly greater (*p* < 0.001) than that in plasma (1.4%), but not significantly different from that in T cells (2.8%). In P01 at enrollment, the pattern agreed with P02 - HIV in monocytes (2.9%) was significantly (*p* < 0.0001) less diverse than either plasma (3.1%) or T-cells (5.0%). At 12 months post-cART, however, the pattern was reversed - HIV in monocytes (4.3%) was significantly (*p* < 0.01) more diverse than both plasma (3.2%) and T-cells (4.1%). Significant differences and variation in the relative levels of genetic diversity suggest distinct evolutionary processes within each peripheral blood compartment.

We performed a spatiotemporal reconstruction of the evolutionary history of tissue and cell populations using a Bayesian phyloanatomic framework [16, 17]. A varying proportion of ancestral lineages were assigned to monocyte origin for all three individuals (Fig 2), with the strongest signal for participant P13. Monocyte viral lineages did not cluster within a monophyletic clade in the time-scaled phylogenies (Fig 2), or the divergence-scaled maximum likelihood (ML) phylogenies (S5 Fig), for any of the three individuals, a finding that does not support a model of completely evolutionarily separate cellular compartments. We confirmed the phyloanatomic inference using quantitative analyses of the pairwise genetic distances between origin-annotated sequences within the ML trees (S1 Table). Because of the relatively sparse sampling of intra-host populations it is not possible to make definitive claims regarding the cellular source of any particular viral population. However, recent models developed in the context of between-host HIV source attribution [43] leverage the topological structure of viral phylogenies to infer that compartment A (e.g.) is likely the source of virus in compartment B if multiple clades of virus from B are nested within the larger clade of virus from A, which can be seen for each of the individual phylogenies (Fig 2).

**Figure 2.**
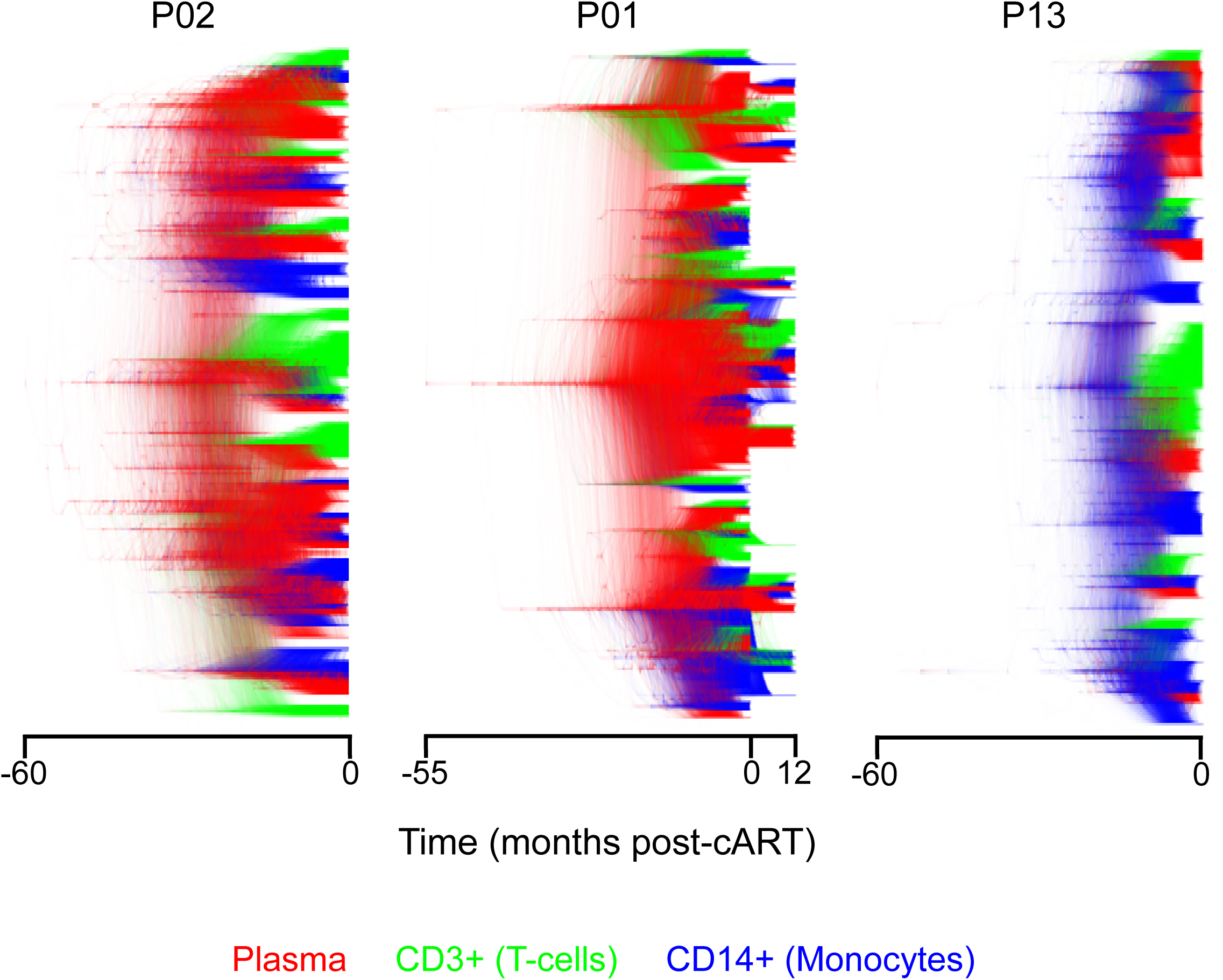
Sampled posterior distribution of Bayesian phylogenetic trees for all HIV *gp120* sequences derived from plasma and sorted peripheral leukocytes in three Thai individuals. A sample (1,000 trees) from the posterior distribution was obtained using an uncorrelated relaxed molecular clock model of evolutionary rate variation across branches [54] and constant population size over time. Branch lengths are scaled in time and colored according to sampling origin (see legend along bottom).

Following tree reconstruction from complete data (Fig 2), sequences were randomly sub-sampled to mitigate compartment sampling bias, and a hierarchical model was incorporated to inform a Bayesian phyloanatomy analysis using all available data in light of the sub-sampling (See Methods). The role of the monocyte ancestral lineages in viral dispersion within the peripheral blood for each of the three individuals was assessed using the Bayesian stochastic search variable selection (BSSVS) diffusion model (Fig 3). We found a consistent signal of significant contribution of infected monocytes to cell-free viral lineages in the plasma, despite differences in sampling time intervals among the individuals, in agreement with earlier studies [13]. Similarly, the contribution of plasma virus to peripheral T-cell lineages was consistently supported (Bayes Factor [BF]>3 [17]). The three participants differed only in the contribution of plasma and peripheral T cells to viral genetic dissemination in monocytes, with contribution from both populations in P02, plasma virus alone in P01, and neither population in P13.

**Figure 3.**
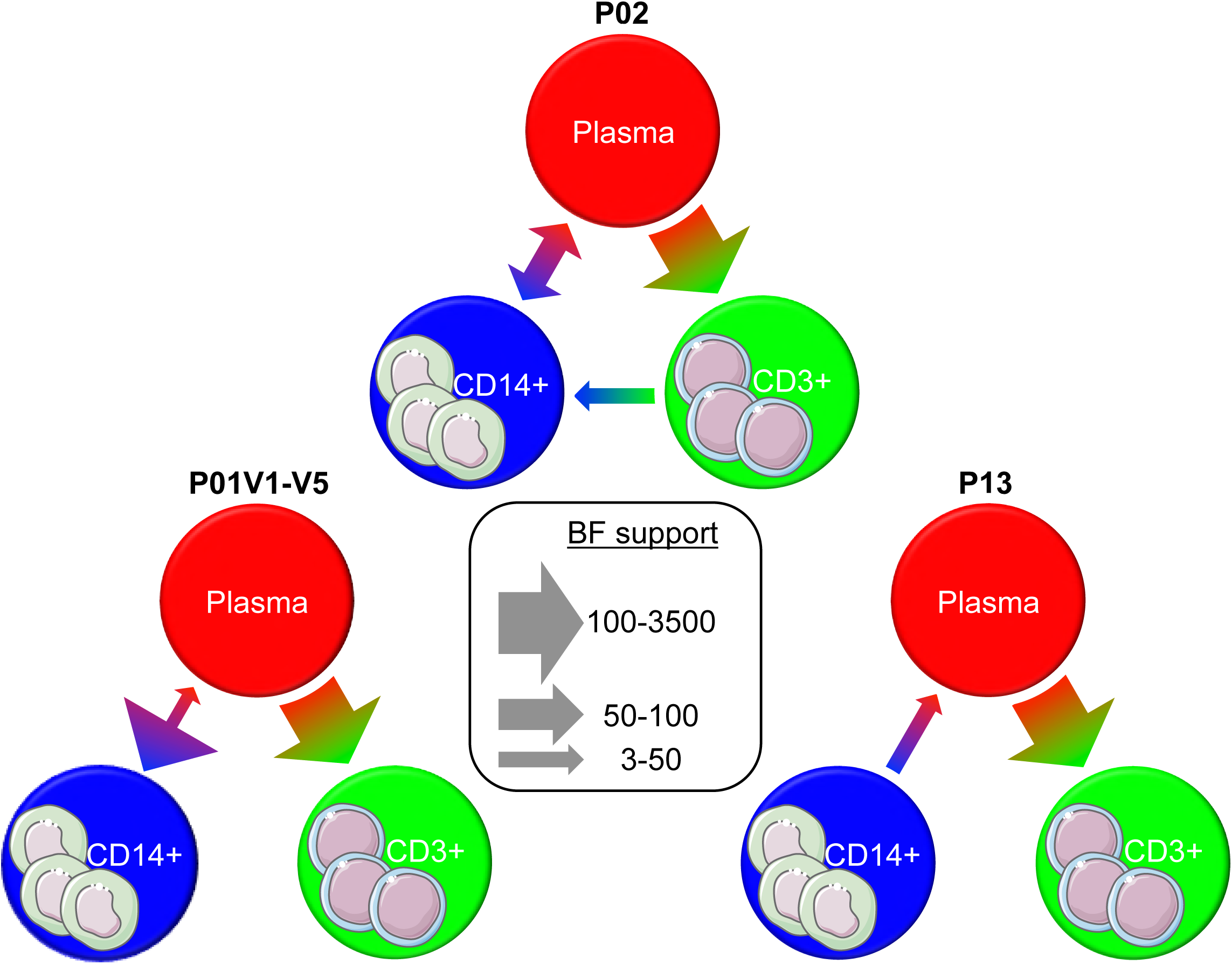
Inferred viral dispersion among peripheral blood compartments in three HIV-1-infected Thai individuals with sufficient samples. The Bayesian stochastic search variable selection (BSSVS) [17] model of asymmetrical transition rates among discrete anatomical locations was utilized in BEAST [49, 61] to reconstruct simultaneously the viral dispersion and *gp120* evolutionary histories for each of the three participants (P01, P02, P13). Transition rates highly supported using (Bayes Factor [lnBF]>3) are depicted as arrows with widths proportional to the BF (legend in center).

Sufficiently large monocyte sequence sample size at the time of enrollment and 12-month follow-up for P01 offered an opportunity for molecular clock dating of specific gene flow events. However, as the patient was being treated during at least a portion of this time, measures were taken to confirm viral divergence from the initial viral sample and, thus, reliable temporal inferences [62, 63]. Analysis of time-dependent phylogenetic clustering of sequences revealed significant genetic distinction between the two different time points (S1 Table), indicating viral population turnover and sufficient evolution during the sampling time interval, despite initiation of therapy. Date randomization tests [51–53] were also performed to determine if greater evolutionary change was occurring over the 12-month period than would be expected by random sampling of available dates, confirming measurable evolution between sampling time points (S8 Fig). Temporal reconstruction in P01 indicated that the median time of transmission for each of the well-supported transitions occurred prior to administration of cART, although extending well into the first year post-therapy in this individual (S7 Fig). Despite potential sampling variation that can accompany the spatial sampling strategy described for these individuals, temporal inferences of the well-supported dispersion patterns in P01 overlapped by as much as 100% among the three replicates of sequence sub-sampling (S7 Fig), indicating the robustness of the molecular clock analysis and BSSVS approach. The results of the phyloanatomy analysis provide evidence for individual-, and potentially, disease-specific (see Conclusions) variation in viral dispersion pathways between peripheral blood targets of HIV, with evidence to suggest their occurrence primarily, but not exclusively, prior to prolonged cART exposure.

We found differences at the amino acid level between virus from plasma, T cells, and monocytes (S9 Fig). These differences were widely dispersed across the GP120 primary sequence, requiring analysis of the rate of change of amino acids at these sites to determine if region-specific evolutionary patterns were responsible for the differences observed for diversity and gene flow among compartments and between individual patients. A population-level approach to site-specific selection detection was developed by adapting the previous dN/dS-based fixed effects likelihood (FEL) model for likelihood ratio testing [44] across population-designated branches (foreground) within the patient-specific ML trees. For P02, thirteen sites were identified as differing significantly (*p* < 0.05) either among any sets of branches or between a specific pair of compartments (i.e., monocytes vs. T cells). These sites were primarily located within the V1-V3 region (Fig 4A). Pairwise differences within this individual were not localized to any particular subregion (e.g., sites differing between T cells and monocytes were distributed across the V2, V3, and V3 regions). The type of selection (positive or negative) also varied according to site. Alternatively, in P13, six sites were identified, primarily located within the 3’ portion of the gene - the C3-V4 region - having no overlap with P02 (Fig 4B). Also in contrast with P02, these sites comprised of differences only between monocytes and remaining locations (i.e., no differences were observed between T cells and plasma), suggesting again a distinct pressure associated with infection of monocytes within this individual. P01 resembled the region-specific pattern of P02, for which significant sites were observed in C1, V2, V3, and C3 regions, three of which were overlapping with sites identified in P02. Only the site located in C3 significantly differed between two of the foreground populations (monocytes and T cells). The results suggest a role for region-specific selection pressures in the distinct gene flow patterns observed in each of the three individuals, with gene-wide selection indicative of a fairly mixed population, but a shift in selection pressure toward the 3’ region indicative of a more restricted gene flow between peripheral blood compartments.

**Figure 4.**
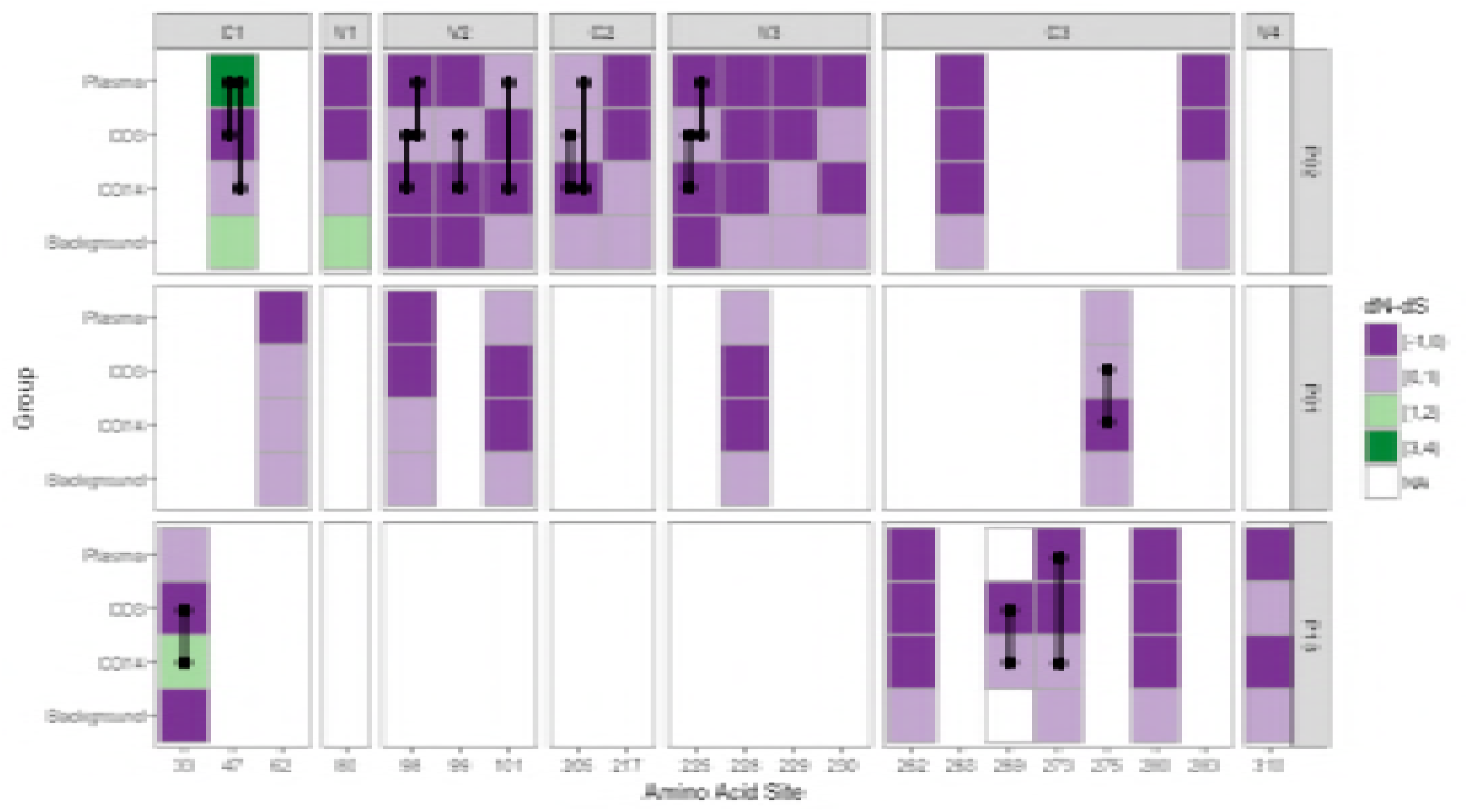
GP120 sites reporting significantly differing selective pressure between branch classifications for each Thai individual. Branches within patient-specific maximum likelihood trees were classified as foreground - plasma, T cells (CD3), or monocytes (CD14) - or background (remaining internal branches). Each population-site pair is colored according to the difference in the rate of non-synonymous (dN) and synonymous (dS) substitutions scaled by the total branch length accompanying site-specific changes. Amino acids comprising constant (C) and variable (V) loops, as defined previously [64], are separated accordingly, with site numbers corresponding to the sequence alignments (aligned between all three patients). Sites reported to differ between any of the three foreground populations are indicated with a black line drawn between the respective population pairs. NA blocks for amino acid site 269 of the P13 sequence alignment indicate no significant overall difference when including background branches. Sites are numbered relative to the beginning of the alignment. P-values ≤ 0.1 were considered significant.

## Discussion

Monocytes harbor a diverse HIV population with potentially many phenotypes [65]. DNA viral populations extracted from monocytes can also be genetically distinct from those isolated from other peripheral blood compartments, such as T cells [11–13, 60]. Expanding on these studies, we investigated genomic RNA, produced via viral replication, thereby mitigating recent concerns about the bias of DNA phylogenetic analysis due to the inclusion of defective viral sequences [66]. Successful isolation of viral genomic RNA and detection of sequence diversity and selective patterns unique to monocyte-derived virus provide sufficient evidence of productive infection of this cell population *in vivo* and a unique role that monocytes play in intra-host HIV evolution. Despite the low prevalence of peripheral monocyte infection *in vivo* [10], their contribution to plasma viral lineages in the three examined patients was significant. On the other hand, the inferred source of monocyte virus varied among individuals, with greater indication of origins outside the peripheral blood (i.e., deep tissue), or at least excluding the present population in the peripheral blood, in two of these patients. Our study furnishes promising evidence of the link between these transmission patterns, viral diversity, and region-specific patterns in selective pressure. Results encourage future investigation into tissue-specific patterns in selection that would shed light onto the potential role of the C3-V4 region in monocyte infectivity. Analyses of longitudinal samples would also be able to reliably distinguish a recent deep-tissue source of transmission from an earlier contributing peripheral blood compartment.

Our finding that diverse tissues potentially act as sources of monocyte infection in specific individuals and time points is not surprising in light of recent evidence of monocyte trafficking [6, 7] and may offer an explanation for the disease statuses of the three Thai individuals. Collectively, the patient diagnoses represented three different stages of HIV-associated neurocognitive disorder (HAND)- P02 as cognitively normal (NL), P01 as mild neurocognitive disorder (MND), and P13 as HIV-associated dementia (HAD) (see Methods for clinical criteria). Though ours is a small sample, the connection is worth noting, as increased monocyte trafficking has been associated with the development of HAND symptoms and of SIV-associated neuropathology in the macaque model of HAND (e.g., [67–70]). The link between a tissue-mediated altered evolutionary landscape and brain infection remains unclear, however, as do the events leading to increased migration.

## Conclusion

Regardless of disease status, the contribution of monocyte infection to ongoing evolution remains an important finding given the inferred timing of initial infection (prior to ART) and previously reported reduced impact of ART on infection of monocytes and macrophages [71]. These cellular compartments could act as an under-appreciated and difficult-to-reach (pharmacologically) HIV reservoir, which, if appropriately targeted by therapy, might improve patient level outcomes and mitigate neuropathological complications of HIV/AIDS.

## Supporting information

**S1 Table. Analysis of compartmentalization of *gp120* sequence data for three HIV-1-infected Thai individuals.**

**S1 Fig. Monocyte cell sorting strategy.** Monocyte/myeloid populations were analyzed by first gating using forward scatter (FSC) and side scatter (SSC) **(A)**. We excluded cell-doublets **(B)** and dead cells **(C)** using FSC height and amine dye, respectively, and then excluded CD3+ T and CD20+ B lymphocytes **(D)**, and CD16+ HLA-DR-NK cells **(E)**. From the HLA-DR+ sub-population **(E)**, three monocyte subsets were distinguished: CD14+CD16*-* classical monocytes, and two subsets of activated monocytes: CD14+CD16+ and CD14lowCD16+ **(F)**. An example of ungated post sort data are shown on the right where the percentage of monocytes population is enriched to 99% **(G)**. The sorted cells show no contamination by CD3+ T and CD20+ B lymphocytes **(H)**, as well as CD16+ HLA-DR-NK cells **(I)**.

**S2 Fig. HIV-1-infected Thai participant diagnosis, sampling time-line, and sequence information.**

**S3 Fig. Linear relationship between estimated genome copy number in monocytes and viral load for all infected Thai participants.** The number of genome copies per million sorted monocytes (y-axis) was estimated in QUALITY [59] using limiting dilution PCR results. The number of cells was determined using FACsorting of CD14 and CD16 molecular markers on 5 mL blood samples. Linear regression analysis of these estimates against viral load (x-axis) was performed in R v3.4.3. Individuals and time points from which sufficient sequence numbers were obtained for Bayesian phylogenetic analysis have been labeled.

**S4 Fig. Viral load in participants with (SGS(+)) and without (SGS(-)) sequenced single genome amplified RNA in monocytes.** Logistic regression analysis of the impact of viral load (y-axis) on the ability to detect and sequence genomic RNA in monocytes of all individuals (x-axis) was performed in R v3.4.3 (p=0.06). Individuals and time points from which sufficient sequence numbers were obtained for Bayesian phylogenetic analysis have been labeled.

**S5 Fig. Maximum likelihood (ML) phylogenies of viral *gp120* sequences for individual HIV-1-infected Thai participants.** ML trees were reconstructed in IQTREE [40] using the best-fit evolutionary model according to Bayesian Information Criterion. Taxa are shaped according to the time of clinical visit in months post-diagnosis (mpd), or post-cART initiation, and are colored according to isolation origin (plasma or sorted peripheral blood mononuclear cell).

**S6 Fig. Viral genetic diversity within plasma and peripheral T-cells and monocytes obtained at specific time points from three individual HIV-1-infected Thai participants.** Viral genetic diversity, represented as the pairwise genetic distances between sequences belonging to the same anatomical compartment, was estimated in R (ape package) using the TN93 evolutionary model [38]. The number of sequences analyzed for each participant-specific compartment is depicted above. Participant (P01, P02, P13) visit number (V1 = 0 months post-cART, V5 = 12 months post-cART) is also indicated. Statistical differences were determined using a non-parametric multiple comparisons test (Dunn package in R) with Bonferroni p-value correction. *p-value<0.05 **p-value<0.01 ***p-value<0.001

**S7 Fig. Inferred timing of viral dispersion among discrete anatomical compartments for participant P01 sample replicates.** Participant *gp120* sequence data were re-sampled thrice (with replacement) according to the minimum number of sequences in one of the three anatomical compartments. The timing, in months post-diagnosis (mpd), of viral dispersion was inferred for individual P01 sample replicates (1-3) using the Bayesian coalescent [50] and phyloanatomy [16] frameworks. *Bayes factor support (>3) indicating a significantly non-zero rate of transition between designated discrete anatomical locations within the Bayesian phylogeny, as determined using the Bayesian stochastic search variable selection model [17] of asymmetric transition rates.

**S8 Fig. Evolutionary rate estimates for true and randomized sampling dates of *gp120* sequence data.** Participant *gp120* sequence data were re-sampled thrice (with replacement) according to the minimum number of sequences in one of the three anatomical compartments, taking into consideration both sampled time points - 0 and 12 months post-cART. Clustered permutation randomization [51, 52] of the true sampling dates (red), was performed in R using the TipDatingBeast package for 5 replicates (Rep, black) [53] and used for Bayesian evolutionary reconstruction in BEAST [49, 50], assuming the uncorrelated, relaxed molecular clock [54] and Bayesian skyline demographic models [50]. Mean evolutionary rate (substitutions/site/month) and 95% high posterior density intervals (error bars) are reported.

**S9 Fig. Amino acid signature pattern analysis for plasma, monocyt and T-cell-associated** gp120 *sequences in three HIV-1-infected Thai individuals at the time of cART initiation*. Amino acid differences are measured as differences in the majority of amino acid present for participant- and anatomical compartment-specific sequence alignments at the time of cART initiation relative to the CM240_AE reference sequence. Differences are denoted by the majority amino acid present for a particular alignment. Amino acids comprising variable loops V1-4, as defined previously [64], are shaded accordingly.

**S10 Fig. Neighbor joining (NJ) phylogeny incorporating all HIV-1-infected Thai individuals for which viral** gp120 *sequences were obtained*. NJ tree reconstruction was performed in MEGA v5.2.1 [35] according to the HKY [36] evolutionary model with gamma-distributed rate variation across sites. Branches are colored according to participants, and those representing HIV-1 subtypes B and AE are outlined.

**S11 Fig. Likelihood mapping of HIV-1-infected Thai participant-specific replicates of uniform sampling from anatomical locations.** Participant *gp120* sequence data were re-sampled thrice (with replacement) according to the minimum number of sequences in one of the three anatomical compartments. Likelihood mapping [47] was performed for each participant (P01, P02, P13) sample replicate (S1-3) in IQTREE [40] using the best-fit evolutionary model according to the Bayesian Information Criterion. Triangular compartments correspond to percent of sequence quartets that were unresolved (center), partially resolved (center edges), or fully resolved (corners) in the phylogenetic tree. Greater than 20% fully resolved quartets was considered sufficient for reliable evolutionary inferences.

## Clinical Trial Protocol

### Additional information

Material has been reviewed by the Walter Reed Army Institute of Research. There is no objection to its presentation and/or publication. The opinions or assertions contained herein are the private views of the author, and are not to be construed as official, or as reflecting true views of the Department of the Army or the Department of Defense. The investigators have adhered to the policies for protection of human subjects as prescribed in AR 70-25. Drs. Ananwonanich and Valcour has served as a consultant to ViiV Healthcare and Merck for consultation unrelated to this study.

The authors report no competing financial interests. Correspondence and requests for materials should be addressed to.

**S1 Table.** Analysis of compartmentalization of *gp120* sequence data for three HIV-1-infected Thai individuals.

| Participant ID | Replicate (Sample) | Time |  | Location |  |
| --- | --- | --- | --- | --- | --- |
|  |  | rb# | r* | rb# | r* |
| P01 | S1 | 0.03 | 0.02 | 0.77 | 0.7 |
|  | S2 | 0.003 | 0.001 | 0.94 | 0.9 |
|  | S3 | 0.02 | 0.009 | 0.4 | 0.36 |
|  | All sequences | 0.08 | 0.005 | 0.26 | 0.4 |
| P02 | S1 | - | - | 0.01 | 0.02 |
|  | S2 | - | - | 0.06 | 0.11 |
|  | S3 | - | - | 0.02 | 0.06 |
|  | All sequences | - | - | 0.35 | 0.19 |
| P13 | S1 | - | - | 0.28 | 0.21 |
|  | S2 | - | - | 0.03 | 0.04 |
|  | S3 | - | - | 0.43 | 0.34 |
|  | All sequences | - | - | 0.52 | 1 |

#correlation coefficient [58] statistic (p-value) representing the number of branches (rb) or \*branch length (r) separating, within the maximum likelihood phylogeny, sequences from separate compartments defined according to sampling time or anatomical sampling origin, or location. Statistical significance was determined using a null distribution of permuted sequences (1,000 permutations). A p-value of $\leq 0.5$ was considered significant.

## Acknowledgments

We thank the study participants and staff from the Thai Red Cross AIDS Research Centre and Pramongkutklao Hospital, Department of Medicine in Bangkok for their valuable contributions to this study. SEARCH is a research collaboration between the Thai Red Cross AIDS Research Centre (TRCARC), the University of Hawaii, and the Department of Retrovirology, U.S. Army Medical Component, Armed Forces Research Institute of Medical Sciences (AFRIMS). This study was supported in part by NIH-NINDS R01 NS053359 (S.R.K.) and NIH-NIGMS U01 GM110749 (S.L.K.P.).

